# Isolation and Characterization of a Novel Alkaline Lipase/Esterase Lip-1420 from a Metagenomic Library

**DOI:** 10.1101/422626

**Authors:** Hee Kyung Lim, Dal Rye Kim, Min Woo Woo, Moon-Sun Hahm, In Taek Hwang

**Affiliations:** Carbon Resources Institute/Green Carbon Catalysis Research Center, Korea Research Institute of Chemical Technology, 141 Gajeong-ro, Yusoeong-gu, Daejeon 34114, Republic of Korea; Bioprogen, VentureTown Jangyoungsil, 407, 105, Sinildong-ro, Daedeok-gu, Daejeon 34324, Korea; Department of Advanced Materials and Chemical Engineering, University of Science & Technology, 217 Gajeong-ro, Yuseong-gu, Daejeon, 34113, Republic of Korea

**Keywords:** Lipase, Esterase, Plant rhizosphere, Soil metagenome, Purification

## Abstract

A novel lipase gene, Lip-1420, was isolated from a metagenomics library constructed from soil samples from reed marshes and from Mt. Jumbong in Korea consisting of 112,500 recombinant plasmids. A selected recombinant plasmid, Lip-1420, was further analyzed which exhibited the maximum lipolytic activity among 18 recombinant fosmids that showed lipolytic activity. Among them, DNA sequence analysis of a 5,513 bp subclone in pUC119 Lip-1420-sub revealed that 11 ORFs were included in the gene according to the blast search from GenBank. The transformant of Lip-1420-ORF3 exhibited lipolytic activity, and plasmid pET21a(+)-Lipase-6H was constructed and transferred to the expression host *E. coli* BL21(DE3). Finally, the lipase Lip1420 was purified from a FPLC system, and the recombinant enzyme was kept in a −70 ºC freezer at a concentration of 1 mg/mL in a buffer solution (50 mM Tris-HCl and 0.15 M NaCl, pH 7.4). The Lip-1420 gene was registered in GenBank (MH628529), and the purified enzyme had an optimal activity at 40 °C and pH 8.0. Kinetic analysis of the Lip-1420 lipase with the substrate *p*-nitrophenyl palmitate was performed at 40 ^°^C and pH 8.0, and the *K*_m_ and *V*_max_ values of the protein were determined to be 0.268 mM and 1.821 units, respectively. The purified Lip-1420 exhibited the maximum enzymatic activity towards *p*-nitrophenyl palmitate, indicating that it is an esterase.

**IMPORTANCE:** This study provides the knowledge to develop a new lipase from the metagenomics library of rhizosphere in Korea using an activity-based method. In addition, the knowledge gained from this study has allowed us to understand that the natural ecosystems are still an unknown genetic material storage report in relation to industrially useful biocatalysts, which are still rich in biodiversity. Moreover, alkaline lipase has great potential for applications in the detergent industry. Especially in the major part of the global industrial enzyme market with high growth potential. It is used, for example, in detergent additives, biopolymer synthesis, and biodiesel production, synthesis of optically pure compounds and food additives as well as in the paper industry, perfume and fragrance blends, biological purification and waste treatment.

## INTRODUCTION

Lipases (EC 3.1.1.3), defined as a carboxylesterase that catalyzes both hydrolysis and the synthesis of long-chain acylglycerols (1), are a group of important industrial biocatalysts that have long been used in the organic chemistry, pharmaceutical, food and leather industries. In addition, they are well-known for their remarkable ability to carry out a wide variety of chemo-, regio- and enantio-selective transformations (2). Lipases have gained much attention worldwide by organic chemists due to their general ease of handling, broad substrate tolerance, high stability towards temperatures and solvents and convenient commercial availability. Recently, lipases have gained great value for producing biodiesel from bio oils by transesterification and esterification of free fatty acids (3).

The discovery of new lipases is continuously required for various industrial developments and will increase the diversity of suitable enzymes for new challenges. Sources of lipase enzymes have been found in nature such as plants, animals and microorganisms (fungi, bacteria, yeast, etc.). The majority of known lipolytic enzymes are of bacterial origin and used in various industries because they are environmentally friendly and non-toxic and have no harmful residues. However, it is well known that only between 0.1 and 1% of microorganisms from an environmental sample can be cultivated using standard techniques, leaving more than 99% of bacterial DNA unexplored (4, 5); thus, searching for new enzymes from microorganisms is limited, and most remain to be exploited. Therefore, metagenomics has become an increasingly important option for discovering new enzymes from complex microbiomes. Metagenomics, defined as the genomic or functional analysis of a bacterial community without its cultivation, represents an interesting technique to discover genes of industrial interest (6). Metagenomics has accelerated the process of discovery of novel biocatalysts by enabling scientists to tap directly into the entire diversity of enzymes held within natural microbial populations (7-10). Lee et al. (10) demonstrated that plant rhizosphere soil is a good resource for screening for novel lipolytic enzymes.

In the lipase industry, in addition to the challenge of discovering new lipase producers, the other challenge is to develop enzymes of excellent quality. For example, the pharmaceutical industry and biodiesel production require stable lipases in water-miscible organic solvents. Halophilic lipases isolated from marine environments are useful in industry (11). Detergent alkaline lipases should have a high activity and stability at a wide range of temperatures and pHs and should be compatible with other components of detergents including metal ions, surfactants and oxidants (12). For instant, they are widely used in the dairy and pharmaceutical industries (13), detergents and surfactants (14), the taste or flavor industry (15), and the agricultural, chemical, cosmetics and perfume industries (14). In particular, the lipase from *Acinetobacter venetianus*, RAG-1, can produce biodiesel using a transesterification process (16). The advantages of producing biodiesel using enzymes are that bio oils can be catalyzed under mild conditions using low energy and that glycerol can be easily recovered from biodiesel (17).

In this report, we describe the isolation, sequence analysis, cloning and expression, transformation and over-expression, enzyme purification and enzymatic characterization of a novel lipase-encoding gene, Lip1420, from a metagenomics library made from a reed marsh at Mt. Jumbong, Korea.

## MATERIALS AND METHODS

### Strains, plasmids, and chemicals

Cultures of the *Escherichia coli* strain EPI-100 were grown on 37 °C Luria-Bertani (LB) agar medium supplemented with the appropriate antibiotics. The concentrations of the antibiotics used for the *E. coli* strains were 50 μg/ml of chloramphenicol, 100 μg/ml of ampicillin, and 50 μg/ml of kanamycin. Plasmids pEpiFOS-5 (Epicenter, Madison, Wis., USA) and pUC119 were used to construct the 112,500 clones for the plant rhizosphere metagenomics library and to sub clone the gene that encoded the lipolytic activity, respectively.

### Construction of the metagenomics library

Soil samples collected from reed marshes at Shinsong-ri, Chungnam and from Mt. Jumbong in the Gangwon province, Korea were used to construct the metagenomics library. After air drying, only the soil was passed through a sieve with a mesh diameter of 1.4 mm. Five grams of the soil samples were dissolved in 15 ml of buffer solution (100 mM Tris-HCl, pH 8.0, Sigma, USA), and then, 100 mM ethylenediaminetetraacetic acid (EDTA, Sigma, USA), 1.5 M sodium chloride (NaCl, Junsei, Japan) and 1% hexadecyl trimethyl ammonium bromide (HTAB, Sigma, USA) were added to the dissolved samples. Then, 100 μl of Proteinase K (100 mg/ml, Sigma, USA) were added to the suspension, followed by shaking at 150 rpm for 30 minutes in a shaking incubator at 37 °C. After the incubation, 1.5 ml of 20% sodium dodecyl sulphate (SDS, Sigma, USA) were added to the reaction mixture and then kept at a constant temperature of 65 °C in a water bath for 30 min. The reaction mixture was centrifuged at 7,000xg for 10 minutes, and the supernatant was decanted. The residue was mixed with a same volume of chloroform/isoamyl alcohol (24: 1) and centrifuged again at 8,000xg for 5 minutes. Only the supernatant containing the DNA was transferred to a new tube. Isopropanol corresponding to 0.6 times the volume of the supernatant was added and mixed, followed by centrifugation at 11,000×g for 5 minutes to obtain a DNA precipitate. The DNA precipitate was washed with 70% ethanol, and then, the ethanol was removed by centrifugation. The resulting precipitate was dried and dissolved in 0.5 ml of a TE (10 mM Tris-HCl and 1 mM EDTA, pH 8.0) buffer solution and stored at 4 °C for library construction. Determination of the amount and size of the isolated DNA was confirmed by pulsed field gel electrophoresis (PFGE). PFGE was performed at 6 V/cm at a 120 degree fixed angle for 6 hours under switch time conditions of 1 to 6 seconds. From this, approximately 1 to 1.5 μg of metagenomic DNA was recovered per 1 g of soil sample, and the size of the isolated DNA was found to be about 20 to 100 kb. The DNA precipitate was electrophoresed on 1% low-melting agarose and eluted with gelase (Epicenter, USA), followed by an end-repair step using an enzyme (Epicenter Biotechnologies, USA). The repaired DNA was transformed into EpiFos (Epicenter, USA) using a library production kit (Epicenter, USA). The transformed *E. coli* was cultured to construct a metagenomics library. To confirm the quality of the library, recombinant plasmids were extracted from randomly selected transformants, treated with restriction enzyme *Bam*HI, and analyzed by electrophoresis. As a result, all the recombinant plasmids were exactly contained insert DNA, and the average size of the inserted metagenome was 35 kb. Soil DNA isolation and DNA purification were performed according to a previously reported method (18, 19). The total 112,500 clones in the metagenomics library in *E. coli* were stored at −80 °C in cryotubes (with more than 500 clones in each pool) until used.

### Lipolytic clone selection and DNA cloning

Lipolytic active clones were selected using LB agar medium which was emulsified with 1% tributyrin as previously described (19). After incubation at 37 °C for 2-4 days, hydrolysis of tributyrin was determined by the clear halo formation around the colonies on LB agar plates. Selected clones were digested with the restriction enzyme *Bam*HI, and the lipolytic active clones were finally selected after electrophoresis. The plasmid DNA was digested with *Pst*I to make several 2-5 kb DNA fragments. The DNA fragments cleaved with *Pst*I were ligated into pUC119 (20) to prepare a secondary shotgun library. The prepared library was plated onto LB agar plates supplemented with ampicillin and tributyrin and incubated at 37 °C for 2 days to isolate lipase active subclones.

### Overexpression and purification of a new lipase Lip-1420

The primers used were 5′- GCGACAGGCCATATGAGTCAGATTCCGGCTGA-3′ and 5′-GCGTCAGGACTCGAGGTGTTTGAGATGTTTGTCG-3′ which add the restriction enzyme recognition site for *Nde*I, and plasmid pET21a(+)-ORF3-6H was amplified by polymerase chain reaction (PCR). The PCR was carried out with 20 μl of 5× PCR buffer solution, 10 μl of N-solution, 2 μl of each primer (100 pmole), 1 μl of template DNA, 1 μl of pfu DNA polymerase and 64 μl of sterilized water for a final volume of 100 μl. Amplification was performed in a thermal Cycler (Bio-Rad, USA) as follows: 30 cycles of 95 ^°^C for 30 sec., 58 ^°^C for 30 sec. and 72 ^°^C for 40 sec. The amplified PCR products were analyzed by 1.5% (w/v) agarose gel electrophoresis and purified using a PCR purification kit (Bioneer Co., South Korea). The fragments were sequenced and introduced into pET21a(+) digested with *Nde*I-*Xho*I, and the resulting plasmid was designated as pET21a(+)-Lip1420-6H. Plasmid pET21a(+)-MBP (or GST) -lipase was generated by overlap extension PCR of the MBP (or GST) gene and lipase gene to improve the solubility of the target protein. Standard recombinant techniques were performed according to the methods of Sambrook et al. (20). DNA sequencing and primer synthesis were performed by Macrogen Inc. (Gasan-dong, Seoul, Republic of Korea). Analysis of the sequence information (nucleotides/amino acids) was performed using the BLAST and ORF finder programs provided by the National Center for Biotechnology Information (NCBI, http://blast.ncbi.nlm.nih.gov).

### Mass production and purification of the new lipase Lip1420

Plasmids were transferred to the expression host *E. coli* BL21(DE3) (F-omp T hsdS_B_(r_B_^-^ m_B_^-^) gal dcm (DE3)) (Stratagene, USA) and plated onto LB plates. A single colony from a fresh plate was picked and grown at 37 °C in 100 ml of Luria-Bertani broth (LB) containing 100 ¼g/ml ampicillin until an OD_600_ of 0.6. They were inoculated in 5000 ml of LB with ampicillin. The cells were grown at 37 °C with shaking until an OD_600_ of 0.6. Expression was induced by adding isopropyl-ɑ-D-thiogalactopyranoside (IPTG, GibcoBRL USA) to a final concentration of 1 mM, and the incubation continued at 20 °C for another 16 h. The cells were harvested by centrifugation at 6,000xg at 4 °C for 10 min. and washed in 10 mL of ice-cold washing buffer (50 mM Tris-HCl and 1 mM EDTA, pH 8.0), which was repeated three times. Then, the cells were resuspended in lysis buffer (50 mM Tris-HCl and 200 mM NaCl, pH 7.0) and disrupted by sonication with a sonic disruptor (CosmoBio Co., LTD). After cell disruption, the cell debris was removed by centrifugation at 12,000×g for 30 min. at 4 °C. The harvested proteins were subjected to SDS-polyacrylamide gel electrophoresis according to the methods of Laemmli (22) and stained with Coomassie Brilliant Blue R250. The lipase Lip1420 was purified from the crude extract with a FPLC system for enzyme harvesting (Bioprogen, Korea). The column chromatography consisted of a Bio-rad HR system, column (2×30 cm, Millipore, USA), Ni-nitrilotriacetic acid resin (Ni-NTA, QIAGEN, Germany), bed volume of 20 ml, flow rate of 4 ml/min, and sample volume of 400 ml. SDS-polyacrylamide gel electrophoresis (SDS-PAGE) was performed according to the methods of Laemmli (22), which consisted of a Hoefer minigel, 15% acrylamide gel with a 1 mm thickness, constant current of 120 V, runtime of 1.5 h, Coomassie Brilliant Blue staining, and sample loading of 20 µl. Resin Ni-NTA was poured into a chromatographic column (bed volume, 20 ml, Millipore, USA), and the column was saturated with nickel ions. A 50 mM solution of metal was applied to the column in the form of sulphate salt to remove unbound metal ions. The column was washed with distilled water and a 50 mM Tris-HCl / 0.15 M NaCl buffer solution (pH 7.4). To elute the proteins, the column was washed with an elution buffer (50 mM Tris-HCl / 0.15 M NaCl (pH 7.4)), and a continuous imidazole gradient was applied by the gradual mixing of a 50 mM Tris-HCl / 0.15 M NaCl buffer (pH 7.4) containing 500 mM imidazole with a 50 mM Tris-HCl / 0.15 M NaCl buffer (pH 7.4). Finally, the mass produced recombinant enzyme was kept in a −70 ºC freezer at a concentration of 1 mg/mL in a buffer solution (50 mM Tris-HCl and 0.15 M NaCl, pH 7.4) stored in separate bottles until used.

### Analysis of the enzyme activity and biochemical characterization

The enzyme activity of the recombinant Lip1420 was measured according to the method mentioned below. The amount of 4-nitrophenol (*p*-NP) released was estimated by absorbance at 410 nm using a *p*-NP standard curve. The optimal temperature and pH condition for the esterase activity of the recombinant protein were examined in 96-well micro plates using various temperatures (ranging from 20 to 70 °C) and pH conditions (ranging from pH 2 to 12). The kinetic parameters (Hanes-Woolf constant *K*_m_ and maximal reaction velocity *V*_max_) were estimated by linear regression from double-reciprocal plots. *p*-Nitrophenyl palmitate (*p*NPP) analysis was measured as the substrate 25 μL of 3 mM *p*-nitrophenyl palmitate (*p*-NPP) solution and 5 μg of purified enzyme and 200 μL of 50 mM Glycine-NaOH buffer (pH8.0) were added to a tube. The final concentration of *p*-NPP was 0.3 mM. The reactants were maintained in a heating block at 40 °C for 10 min. and then measured by absorbance at 410 nm with a spectrophotometer. The activity was calculated by substituting the measured values into a *p*-Nitrophenol (*p*-NP) standard curve. One unit (U) of esterase activity is defined as one μmol of *p*-NP released per mg protein min^-1^ (23).

## RESULTS

### Construction of the rhizosphere soil metagenomics libraries

We constructed three metagenomics libraries in a fosmid using DNA extracted from plant rhizosphere soil samples collected from three different locations in the Republic of Korea. The number of clones from the three libraries was as follows: 52,500 clones from the soils of Mt. Jumbong in Gangwon-do and 60,000 clones from the rhizosphere soils of reed plants in Sinseong-ri, Hansan-myeon, Seocheon-gun, Chungcheongnam-do. Altogether, 112,500 metagenomic clones were obtained with an average DNA insert size of approximately 35-40 kb when randomly selected clones were analyzed by agarose gel electrophoresis after *Bam*HI digestion (data not shown), covering at least 5 Gb of rhizosphere microbial genomes.

### Selection of a metagenomic gene encoding lipase Lip-1420

The plant rhizosphere soil metagenomics libraries were screened to find clones that exhibited lipolysis activity. Forty-six clones with lipolytic activities were selected from the prepared 112,500 soil metagenomics library using an activity-based screening method. The *Bam*HI restriction enzyme was used to remove duplicate clones and to isolate 18 unique genes that hydrolyzed tributyrin. Among the 18 unique genes, the Lip-1420 clone was selected, and Lip-1420-sub was constructed using the *Pst*I restriction enzyme (Fig. 1). Lip-1420-sub clones, which contained the smallest DNA insert and showed the best lipolytic activity, were selected. The Lip-1420-sub was completely sequenced and was found to carry a 5,513-bp DNA fragment as an insert. As a result of analyzing the information of the subclone Lip-1420, the characteristics and function of each ORF were summarized and evaluated (Table 1). The genome sequence of the Lip-1420-sub 5,513-bp DNA fragment was found to contain more than 11 open reading frames (ORFs). Of these, ORF1-4 was selected and transformed. The others were excluded because they were too short to be included as a candidate. The information of ORF1 showed a 48% homology with a protein of unknown function in *Thiocapsa roseopersicina* and consisted of 1,590 bp (529 amino acids). ORF2 showed a 50% homology with LLM class flavin-dependent oxidoreductase of *Blastopirellula marina* and consisted of 1,236 bp (411 amino acids). ORF3 showed a 78% homology with an esterase/lipase protein of an *uncultured organism* and consisted of 885 bp (294 amino acids). ORF4 showed a 37% homology with the alpha/beta hydrolase of *Pseudorhodoplanes sinuspersici* and consisted of 825 bp (274 amino acids). For transformation of these 4 ORFs, primers were prepared (Table 2), and transformation into *E. coli* was done for each one. However, only the Lip-1420-ORF3 transformants were further cultured to mass-produce lipase Lip-1420 because only ORF-3 showed the expected lipase activity.

**Fig. 1.**
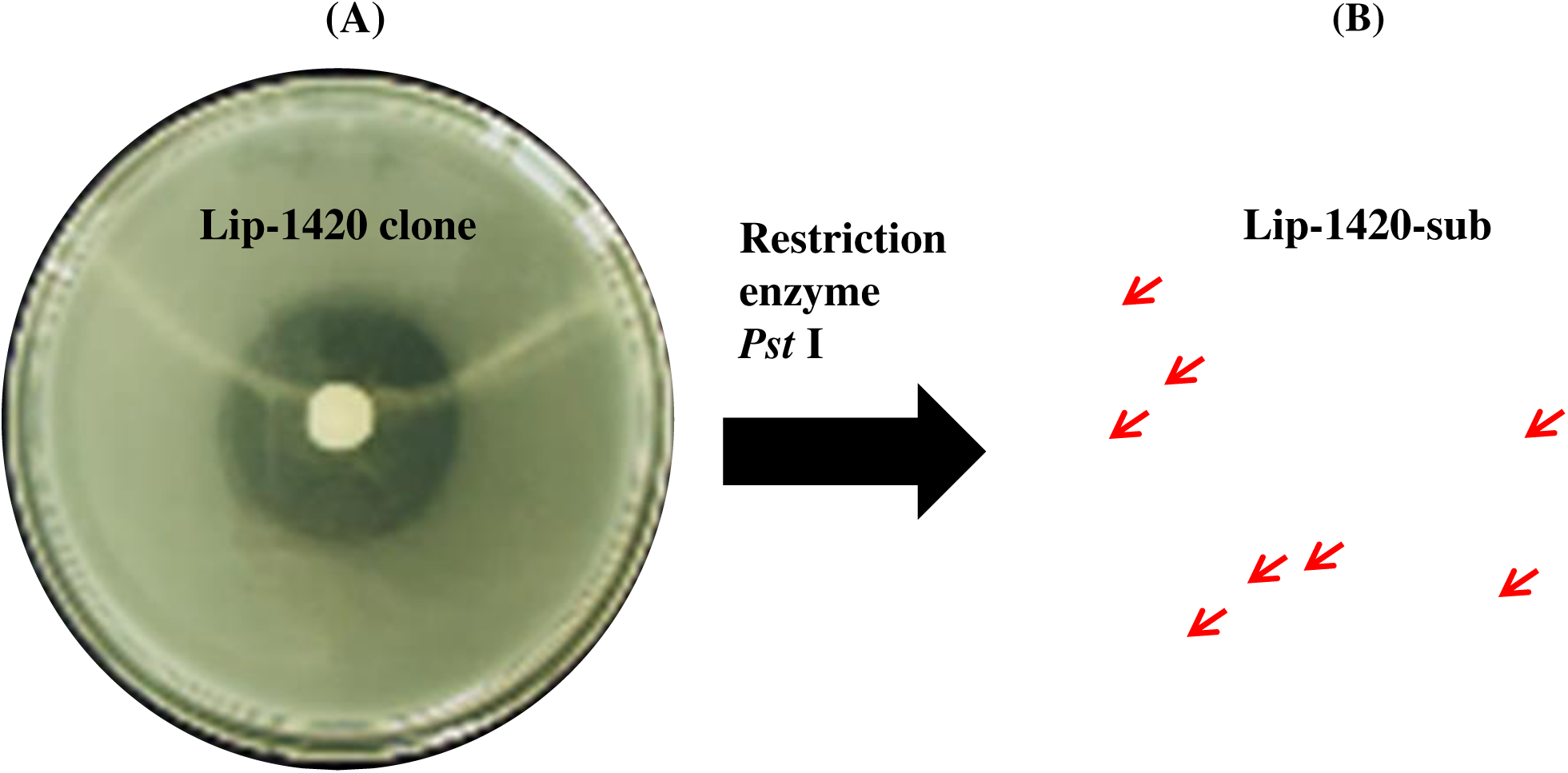
**The Lip-1420 lipase clone selected from 112,500 metagenomic clones (A). Lip-1420-sub gene was selected from restriction enzyme *Pst*I treatment (B).** Agar plate contain tributyrin, and showed clear zone around clones. The larger clear zone means the greater activity.

**Table 1.** **Information of Lip-1420-sub gene consisting open reading frames (ORF) from GeneBank**

| ORF | Length (bp/aa) | Description | Strain (ID %) | Accession |
| --- | --- | --- | --- | --- |
| ORF1 | 1590/529 | Protein of unknown function | *Thiocapsa roseopersicina* (48%) | SDW97003.1 |
| ORF2 | 1236/411 | LLM class flavin-dependent oxidoreductase | *Blastopirellula marina* (50%) | WP_002655372.1 |
| ORF3 |  |  |  |  |
| ORF4 | 825/274 | alpha/beta hydrolase | *Pseudorhodoplanes sinuspersici* (37%) | WP_086087576.1 |
| ORF5 | 495/164 | PREDICTED: 3-hydroxyisobutyryl-CoA hydrolase, mitochondrial-like | *Polistes dominula* (26%) | XP_015178720.1 |
| ORF6 | 435/144 | 4-methyl-5-beta-hydroxyethyl thiazole kinase | *Listeria welshimeri* serovar 6b str. SLCC5334(35%) | A0AFC3.1 |
| ORF7 | 423/140 | No significant similarity found |  |  |
| ORF8 | 402/133 | RNA-directed RNA polymerase | *Pseudomonas virus phi6*(42%) | P11124.3 |
| ORF9 | 327/108 | Phosphoribosylformylglycinamidine synthase subunit PurL | *Leifsonia xyli* subsp. xyli str. CTCB07(31%) | Q6ACC3.1 |
| ORF10 | 294/97 | PEP12 homolog | *Arabidopsis thaliana* (67%) | Q39233.1 |
| ORF11 | 237/78 | No significant similarity found |  |  |

**Table 2.**
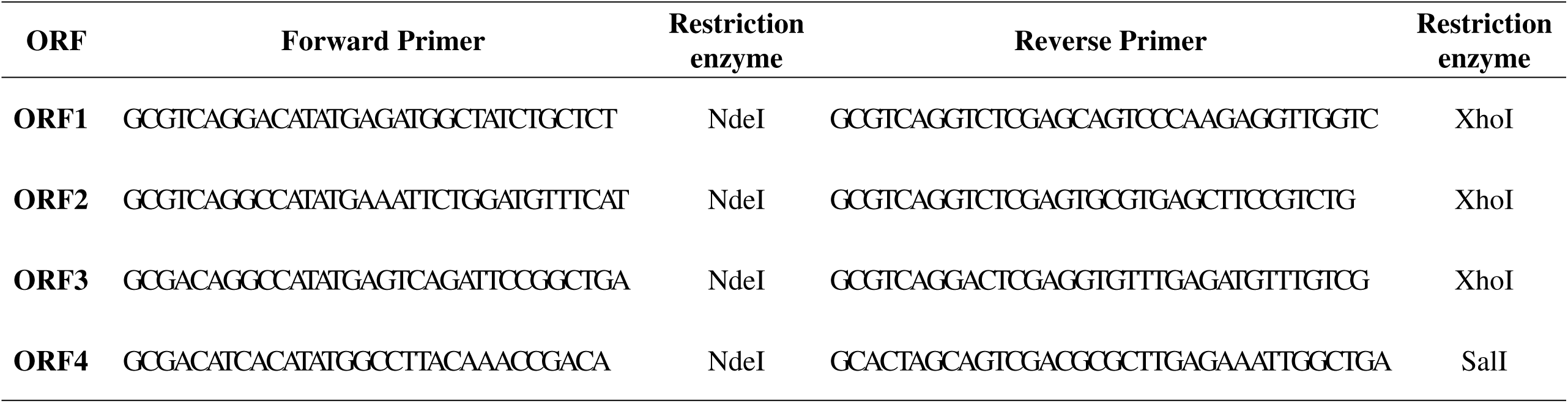
**Primer construction for Lip-1420 gene transformation into *E. coli*.**

### Mass production and purification of the new lipase Lip1420

For the mass production of Lip-1420, pET21a+ORF3-6H (Fig. 2) was used as a basic vector construct, and transformants of the *E. coli* BL21 (DE3) and BL21 (DE3)RIL strains were compared. The expressed protein size was 33.5 kDa (906 bp; 302 a.a.) and was more well expressed in *E. coli* BL21(DE3) than in the variant BL21(DE3)RIL. The Lip-1420 lipase was expressed as the insoluble and soluble forms at 60% and 40% (Fig. 3A), respectively. Immobilized metal affinity chromatography was used for the purification of the enzyme, and Ni-nitrilotriacetic acid (Ni-NTA, QIAGEN, Germany) was used as the resin (Fig. 3B). The final preparation of the Lip-1420 lipase gave a major single band on SDS-PAGE with a molecular weight (MW) of 33.5 kDa, which appeared just below the 35 kDa MW marker. Finally, we harvested a total volume of 400 mL (10 mL × 40 bottles, 1 mg/mL) of Lip-1420 protein with a Bio-Rad FPLC system (Fig. 4A) and subjected it to SDS-polyacrylamide gel electrophoresis according to the collection number and kept the protein in a −70 °C freezer at 1 mg/ml in buffer solution (50 mM Tris-HCl and 0.15 M NaCl, pH 7.4) (Fig. 4B).

**Fig. 2.**
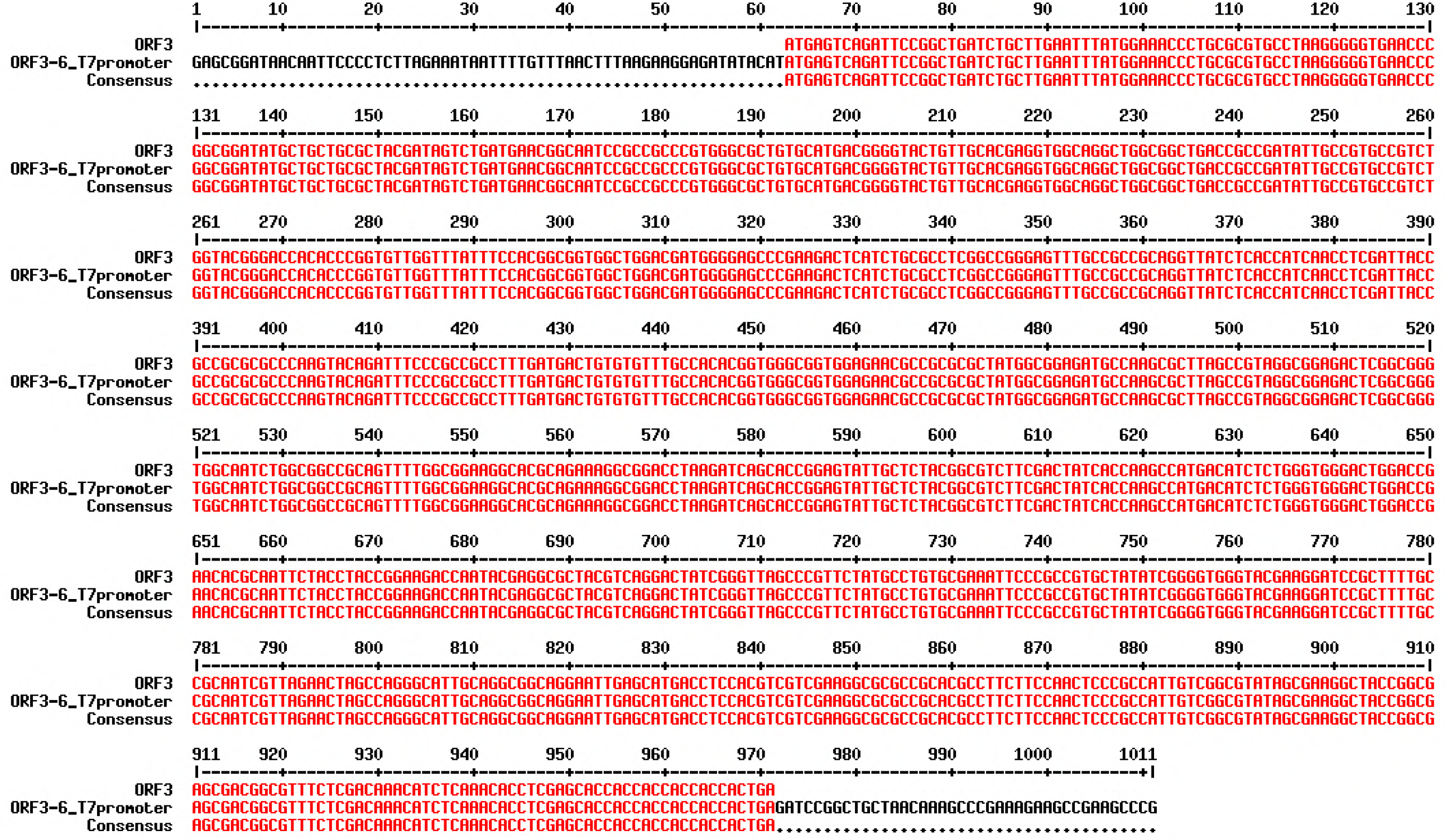
**Sequencing comparision for pET21a+ORF3-6H expression using T7 promoter.**

**Fig. 3.**
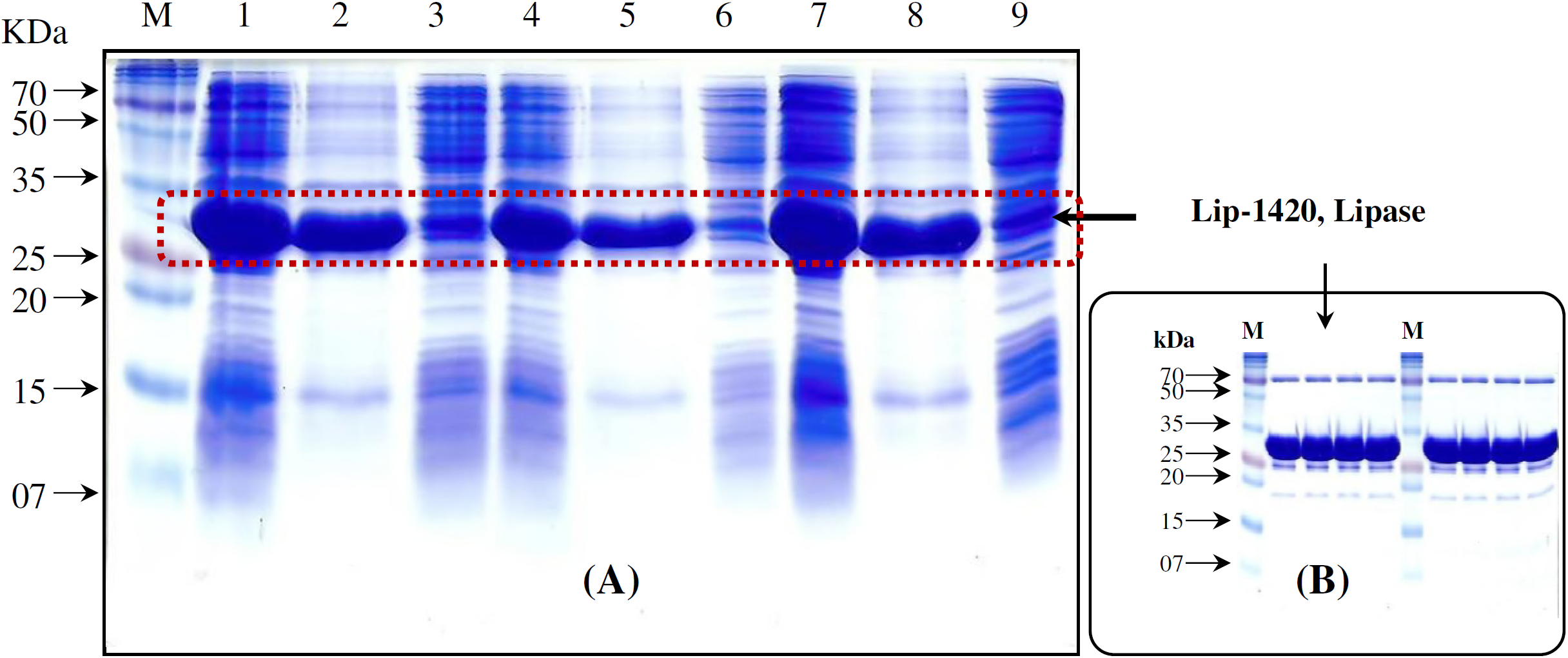
**SDS Page of the recombinant lipase in pET21a+ORF3-6H expression vector (A) and the purified enzyme with immobilized metal affinity column chromatography (B).** M, Marker (ELPIS); 1, total fraction; 2, insoluble fraction; 3, soluble fraction; 4, total fraction (2 times dilution); 5, insoluble fraction (2 times dilution); 6, soluble fraction (2 times dilution); 7, total fraction; 8, insoluble fraction; 9, soluble fraction.

**Fig. 4.**
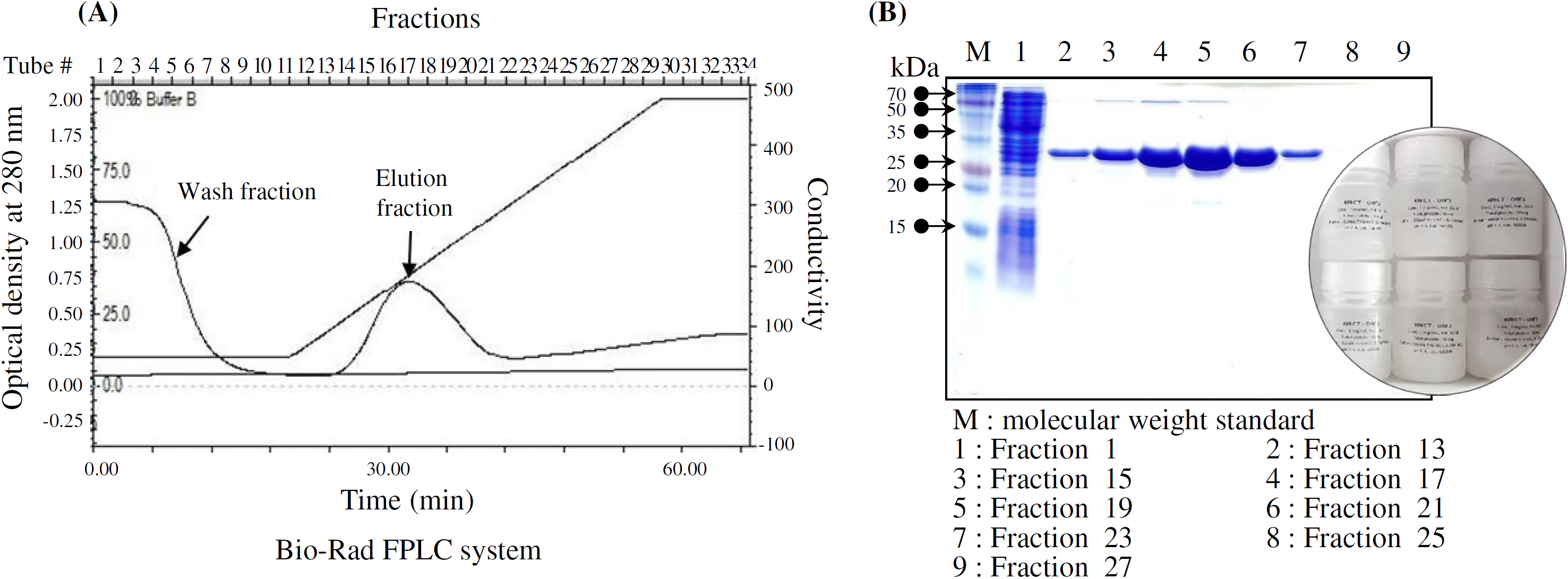
**FPLC system for enzyme harvesting (A) and SDS-PAGE analysis (B).** Column chromatography: Bio-rad HR system, Column (2×30 cm, Millipore, USA), Resin (Ni-NTA, QIAGEN, Germany), Bed volume 20 ml, Flow rate 4 ml/min, Sample 400 ml; SDS-PAGE analysis system: Hoefer minigel, Acrylamide gel 15%, 1 mm thickness, 120V constant current, 1.5 hr, Coomassie Brilliant Blue staining, Sample loading 20µl. The final lipase production was 1 mg/ml in buffer solution (50 mM Tris-HCl, 0.15 M NaCl, pH 7.4).

### Lipolysis activity of the Lip-1420 lipase

Using affinity chromatography, we purified the Lip-1420 lipase with a molecular weight of 33.5 kDa and investigated its enzyme characteristics. The final purified enzyme, Lip-1420, was identified in agar medium supplemented with tributyrin (C_4_), which is the actual enzyme for the lipolysis activity expressed by the metagenomic subclone (Fig. 1). Lip-1420 hydrolyzed a triglyceride composed of a short chain tributyrin (C_4_) but did not show activity for C_10_, C_16_, and C_18_ long chain triglycerides.

### Optimal conditions and enzymatic kinetics of the Lip-1420 lipase

The biochemical properties of the purified recombinant enzyme were examined under optimal conditions. This enzyme has an optimum pH of 8.0, and its activity was maintained at 80% at pH 7.0 or 9.0 and decreased at pH 6.0 or 10.0 (Fig. 5 A), suggesting that it is an alkaline enzyme. This enzyme was active between 30 and 50 °C; however, 40 °C was the optimal temperature for its activity (Fig. 5 B), and it was stable for 24 h at the optimum temperature of 40 ^°^C; however, the activity decreased at 30 or 60 ^°^C (data was not shown). Kinetic analysis of the Lip-1420 with the substrate *p*-nitrophenyl palmitate was performed at 40 ^°^C and pH 8.0. The *K*_m_ and *V*_max_ values of the protein with *p*-nitrophenyl palmitate as a substrate were 0.268 μM and 75.758 units, respectively (Fig. 6). Generally, the relative enzymatic activity gradually decreased against the longer chain length substrate of the *p*-nitrophenyl ester of fatty acids.

**Fig. 5.**
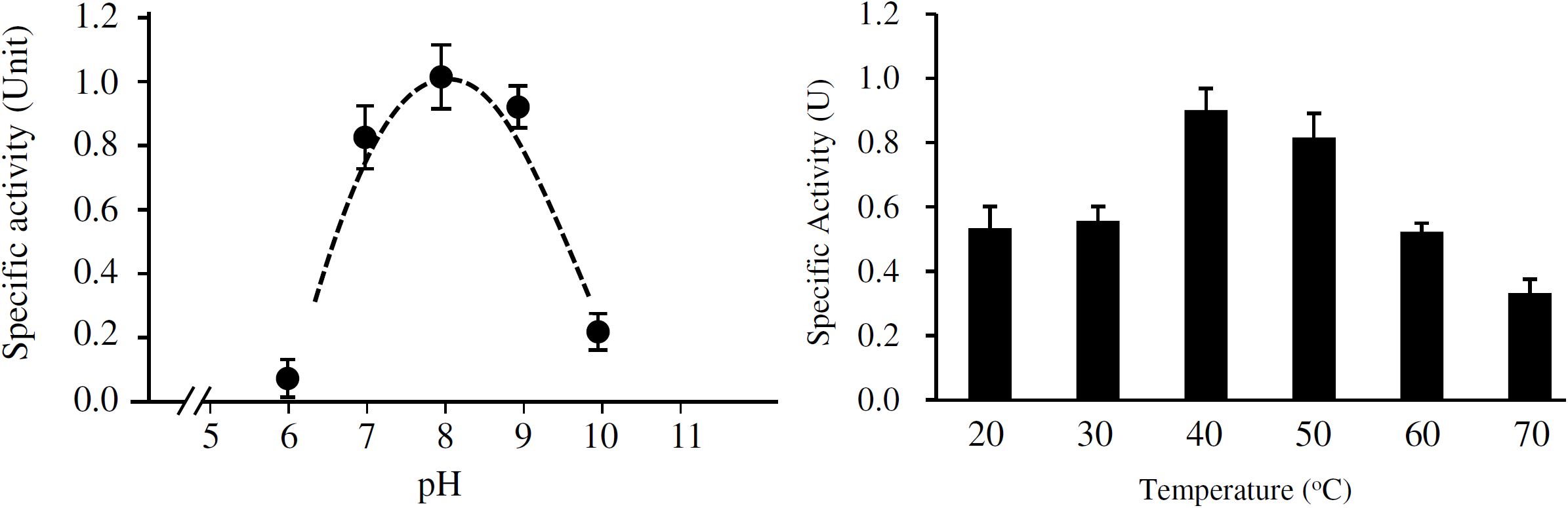
**Optimal condition of the enzyme activity. (A), pH; (B), temperature.** The enzyme activity was assayed at 50 ◦C in 50 mM citric buffer (pH 2∼6.5), phosphate buffer (pH 7∼9), and glycine buffer (pH 9.5∼12).

**Fig. 6.**
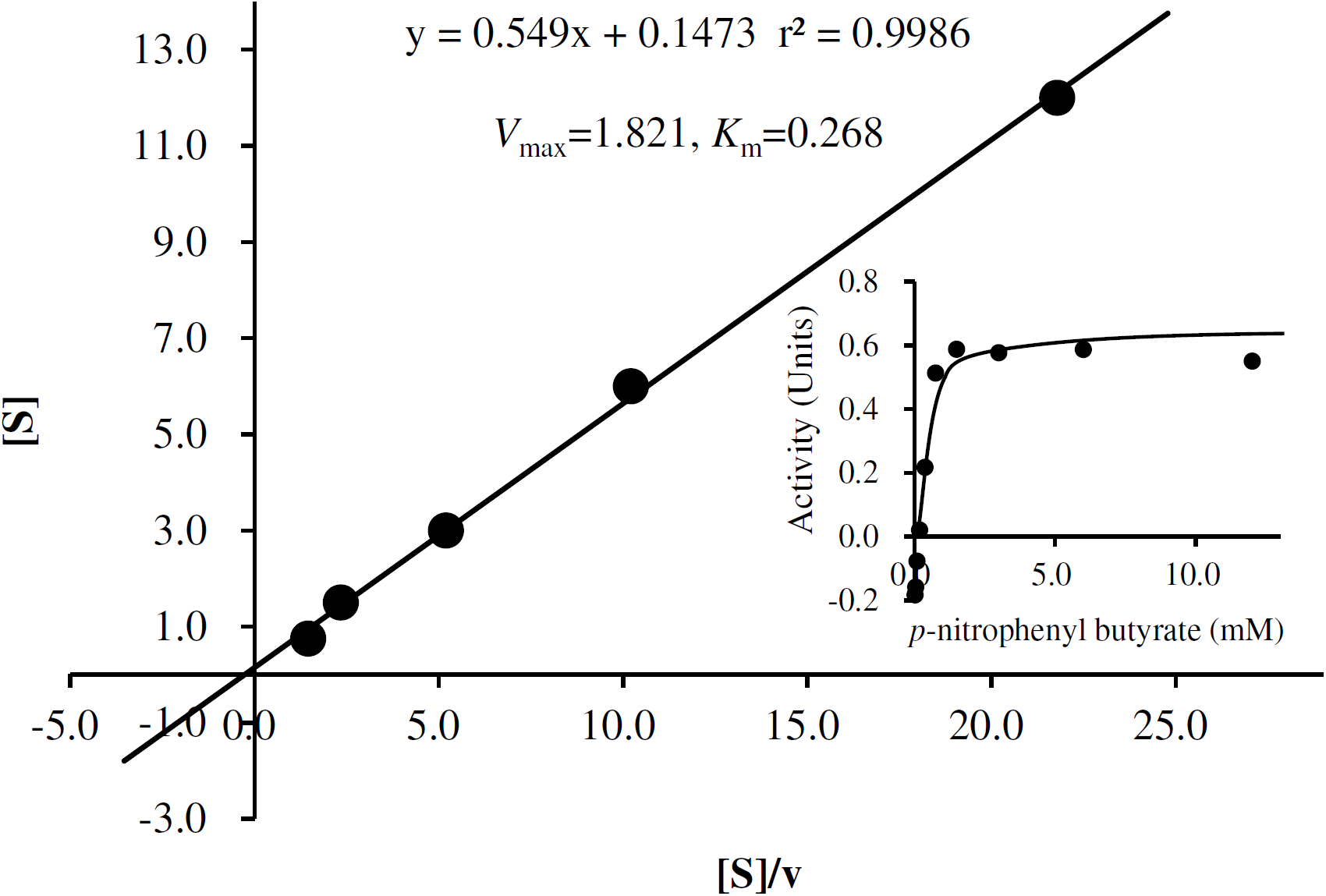
**Lineweaver-Burk double reciprocal analysis of Lip-1420 lipase/esterase activity with *p*-nitrophenyl palmitate as the substrate.** 1 unit of releasing 1 μmol of free *p*-nitrophenol per minute.

## DISCUSSIONS

To use an enzyme industrially, first, a gene encoding a functional enzyme has to be found, and then, its effect has to be enhanced, and finally, it can be mass-produced. Researchers generally collect new variants to discover new functional genes. Many new genes and enzymes can be extracted from unculturable microbes in various habitats using a metagenomics approach (24). However, the use of specific environmental samples to construct metagenomics libraries has been considered to increase the frequency of finding desirable genes in a metagenomics library using functional-centered analysis (25). Because lipids are the main component of the plant root epidermis, lipids or lipid derivatives liberated from plant root boundaries may be nutrients or important signaling molecules in microorganisms near the root (26, 27). Because various lipolytic enzyme-producing bacteria use plant rhizosphere lipids, metagenomics libraries made from plant rhizosphere ecosystems can be a rich source of new lipolysis enzymes.

In this study, a metagenomic library with 112,500 clones was constructed using the rootstock soil of reed marshes and soil samples from Jumbong Mountain in Korea. Thus, 18 new lipolytic clones were isolated after eliminating duplication from the 46 lipolytic clones selected primarily by using tributyrin as a substrate. Among these new clones, the Lip-1420 clone exhibited the best lipolysis activity, and the Lip-1420-subclone was constructed by ligating into the *Pst*I restriction site. According to the analysis of the Lip-1420-subclone gene information using a GenBank blast search, the Lip-1420-subclone had more than 11 ORFs. The Lip-1420-sub-ORF3 transformant was the only one to show a lipolysis activity and revealed a novel gene (GenBank Access No. MH628529). The gene was expressed in *E. coli* BL21(DE3), and 60 mg of purified protein were produced with a Bio-Rad FPLC system using immobilized metal affinity chromatography from a 5 L culture of an *E. coli* transformant. The protein was kept in a −70 °C freezer (1 mg/ml, 10 ml × 6 bottles). Finally, the biochemical characteristics of the enzyme was analyzed. Lip-1420 hydrolyzed a short chain triglyceride tributyrin but cannot digest long-chain triglycerides more than C_10_. Enzymatic lipolysis activity can be detected when a substrate is dissolved in water. However, more than C_12_ triglycerides could not be used as substrates for the analysis of lipolysis activity with the agar plate method without the addition of a surfactant because these are insoluble in water. However, *p*-nitrophenyl palmitate (C_18_) could be used as a substrate to evaluate the esterase activity after it is dissolved in propanol. The optimal temperature and pH of this enzyme was found to be 40 ^°^C and a pH of 8.0, respectively. Under the optimal conditions, the *K*_m_ and *V*_max_ values of the enzyme with *p*-nitrophenyl palmitate as the substrate were 0.268 μM and 75.758 units, respectively.

Lipases/esterases are now a major part of the global industrial enzyme market with high growth potential. They are used as a detergent additive, in biopolymer synthesis, in biodiesel production, in the synthesis of optically pure compounds and as food additives (remodeling of fat to develop sensory and nutritional quality) as well as in the paper industry (paper pulp removal) and as anti-inflammatory agents and in cosmetics (fragrances and fragrance mixtures) and pesticides (herbicides and pesticides) and in biological purification and waste disposal (28-31).

In conclusion, a novel alkaline lipase Lip-1420 was discovered using an activity-based method from a metagenomics library of rhizosphere in Korea. Lip-1420 was active and stable in a broad alkaline pH range (pH 7.0 to 9.0). Lipases that are active and stable in alkaline media have great potential for applications in the detergent industry. For example, lipases that will work under alkaline conditions as fat stain removers are desirable (32). Industrial ecosystems are still biodiversity-rich, and there are still yet unknown reservoirs of genetic material with respect to industrially useful biocatalysts (33, 34). Because the pH instability of enzymes is one of the major bottlenecks in expanding the scope of enzyme utilization, the stability mechanism of acidic and alkaline enzymes is very important and interesting both academically as well as industrially. This new lipase Lip-1420 could be further used to achieve a better understanding of enzyme mechanisms and structure-function relationships.

## ACKNOWLEDGEMENT

This research was supported by a grant (NRF-2017M1A2A2049104) funded by the Ministry of Education, Republic of Korea.

## REFERENCES

1. Gupta, R., N. Gupta, and P. Rathi. 2004. Bacterial lipases: an overview of production, purification and biochemical properties. Appl. Mocrobiol. Biotechnol. 64:763–781.

2. Kumar, A., K. Dhar, S.S. Kanwar, and P.K. Arora. 2016. Lipase catalysis in organic solvents: advantages and applications. Biological Procedures Online 18(2):1–11.

3. Bajaj, A., P. Lohan, P.N. Jha, and R. Mehrotra. 2010. Biodiesel production through lipase catalyzed transesterification: an overview. J. Mol. Catal. B Enzyme 62:9–14.

4. Torsvik, V., J. Goksoyr, and F.L. Daae. 1990. High diversity in DNA of soil bacteria. Appl. Environ. Microbiol. 56:782–787.

5. Steele, H.L., K-E Jaeger, R. Daniel, and W.R. Streit. 2009. A dvances in Recovery of Novel Biocatalysts from Metagenomes. J. Mol. Microbiol. Biotechnol. 16:25–37.

6. Riesenfeld, C.S., P.D. Schloss, and J. Handelsman. 2004. Metagenomics: genomic analysis of microbial communities. Annu. Rev. Genet 38:525–552.

7. Su, J., F. Zhang, W. Sun, V. Karuppiah, and G. Zhang. 2015. A new alkaline lipase obtained from the metagenome of marine sponge *Ircinia* sp. World J. Microbiol. Biotechnol. 31:1093–1102.

8. Ranjan, R., A. Grover, R.K. Kapardar, and R. Sharma. 2005. Isolation of novel lipolytic genes from uncultured bacteria of pond water. Biochem. Bioph. Res. Co. 335:57–65.

9. Glogauer, A., V. Martini, H. Faoro, G. Couto, M. Muller-Santos, R. Monteiro, D. Mitchell, E. Souza, F. Pedrosa, and N. Kireger. 2011. Identification and characterization of a new true lipase isolated through metagenomic approach. Microb. Cell. Fact. 10:54–69.

10. Lee, M.H., K.S. Hong, S. Malhotra, J-H Park, E.C. Hwang, H.K. Choi, Y.S. Kim, W. Tao, and S-W. Lee. 2010. A new esterase EstD2 isolated from plant rhizosphere soil metagenome. Appl. Microbiol. Biotechnol. 88:1125–1134.

11. Selvin, J., J. Kennedy, D. Lejon, G. Kiran, and A. Dobson. 2012. Isolation identification and biochemical characterization of a novel halo-tolerant lipase from the metagenome of the marine sponge *Haliclona simulans*. Microb. Cell Fact. 11:72–86.

12. Wang, Y., K.C. Srivastava, G-J Shen, and H.Y. Wang. 1995. Thermostable alkaline lipase from a newly isolated thermophilic *Bacillus*, strain A30-1 (ATCC 53841). J. Ferment. Bioeng. 79:433–438.

13. Hoshino, T., T. Sasaki, Y. Watanabe, T. Nagasawa, and T. Yamane. 1992. Purification and some characteristics of extracellular lipase from *Fusarium oxysporum* f. sp. *Lini*,” Bioscience Biotechnology and Biochemistry 56:660–664.

14. Hasan, F., A. Sanh, and A. Hameed. 2005. “Industrial applications of microbial lipases,” Enzyme and Microbial Technology 39:235–251.

15. Ochoa, L.C., C.R. Gomez, G.V. Alfaro, and R. Ros. 2005. “Screening, purification and characterization of the thermoalkalophilic lipase produced by bacillus *thermoleovorans* CCR11” Enzyme and Microbial Technology 37:648–654.

16. Ericks, S.A., and C.R. Rita. 2008. “Transesterification activity of a novel lipase from acinetobacter venetianus RAG-1,” Antonie van Leeuwenhoek 94: 621-625.

17. Bajaj, A., P. Lohan, P. Jha, and R. Mehrotra. 2010. “Biodiesel production through lipase catalyzed transesterification,” J. of Molecular Catalysis B: Enzymatic 62:9–14.

18. Rondon, M.R., P.R. August, A.D. Bettermann, S.F. Brady, T.H. Grossman, M.R. Liles, K.A. Loiacono, B.A. Lynch, I.A. MacNeil, C. Minor, C.L. Tiong, M. Gilman, M.S. Osburne, J. Clardy, J. Handelsman, and R.M. Goodman. 2000. Cloning the soil metagenome: a strategy for accessing the genetic and functional diversity of uncultured microorganisms. Appl. Environ. Microbiol. 66:2541–2547.

19. Lee, S.W., K. Won, H.K. Lim, J.C. Kim, G.J. Choi, and K.Y. Cho. 2004. Screening for novel lipolytic enzymes from uncultured soil microorganisms. Appl. Microbiol. Biotechnol. 65:720–726.

20. Vieira, J., and J. Messing. 1987. Production of single-stranded plasmid DNA. Methods in Enzymology 153:3–11.

21. Sambrook, J., E.F. Fritsch, and T. Maniatis. 1989. Molecular cloning. A laboratory manual, 2nd ed. Cold Spring Harbor Laboratory Press, New York.

22. Laemmli, U.K. 1970. Cleavage of Structural Proteins during the Assembly of the Head of Bacteriophage T4. Nature 227:680–685

23. Hoshino, T., T. Sasaki, Y. Watanabe, T. Nagasawa, and T. Yamane. 1992. Purification and some characteristics of extracellular lipase from *Fusarium oxysporum* f. sp. *Lini*. Biosci. Biotech. & Biochem. 56:660–664.

24. Handelsman, J. 2004. Metagenomics: application of genomics to uncultured microorganisms. Microbiol. Mol. Biol. Rev. 68(4):669–85.

25. Simon, C., and R. Daniel. 2009. Achievements and new knowledge unraveled by metagenomic approaches. Applied Microbiology and Biotechnology 85(2):265–276.

26. Sandkvist, M. 2001. Biology of type II secretion. Mol. Microbiol. 40(2):271–83.

27. Wang, X. 2004. Lipid signaling. Curr. Opin. Plant Biol. 7:329–336.

28. Haki, G.D., and S.K. Rakshit. 2003. Developments in industrially important thermostable enzymes: a review. Bioresour. Technol. 89(1):17–34.

29. Joseph, B., P.W. Ramteke, G. Thomas, N. Shrivastava. 2007. Standard review cold-active microbial lipases: a versatile tool for industrial applications. Biotechnol. Mol. Biol. Rev. 2(2):39–48.

30. López-López, O., M.E. Cerdán, and M.I.G. Siso. 2014. New Extremophilic Lipases and Esterases from Metagenomics. Current Protein and Peptide Science 15:445–455.

31. Bajaj, A., P. Lohan, P.N. Jha, and R. Mehrotra. 2010. Biodiesel production through lipase catalyzed transesterification: An overview. J. of Molecular Catalysis B: Enzymatic 62:9–14.

32. Cherif, S., S. Mnif, F. Hadrich, S. Abdelkafi, and S. Sayadi. 2011. A newly high alkaline lipase: an ideal choice for application in detergent formulations. Lipids Health Dis. 10:221–229.

33. Lailaja, V.P., and M. Chandrasekaran. 2013. Detergent compatible alkaline lipase produced by marine *Bacillus smithii* BTMS 11. World J Microbiol Biotechnol 29(8):1349–60.

34. Su, J., F. Zhang, W. Sun, V. Karuppiah, G. Zhang, Z. Li, and Q. Jiang. 2015. A new alkaline lipase obtained from the metagenome of marine sponge *Ircinia* sp. World J. Microbiol. Biotechnol. 31:1093–1102.

